# Workplace Harassment and Risk of Depression Among Adults in the United States

**DOI:** 10.1101/618355

**Authors:** Livingstone Aduse-Poku, Shayester Jahanfar

## Abstract

**Purpose:** The purpose of the study is to perform an analysis of the relationship between workplace harassment and risk depression among adults in the United States.

**Methods:** Across-sectional study of 33772 adults was done using the National Health Interview Survey (NHIS) sample adult data in the USA. The variables collected included depression, harassment, age, race, sex, and marital status. Data was analyzed using IBM Statistical Package for the Social Sciences (SPSS) for windows, version 25.0.

**Results:** Results showed a possible relationship between workplace harassment and depression among adults. Adults who were harassed were 0.39 times more likely to be depressed than those who were not harassed (95%CI: 0.45-0.81), after adjusting for sex, age, race, marital status, pay, and number of years on the job, the odd was almost the same [AOR (95%CI):0.55(0.40-0.77)].

**Conclusion:** The findings of the study show that there is a positive association between workplace harassment and depression.

## INTRODUCTION

Workplace harassment is an unwelcome repeated conduct based on race, color, religion, sex, age or disability targeted at an individual or group of individuals at the workplace (United States Equal Employment Opportunity Commission, 2018). Despite decades of research and efforts to lower its incidence, workplace harassment is still on the rise (Raver, Nishii, Kozlowski & Steve, 2010). The United State Equal Employment Opportunity Commission (USEEOC) in 2006, reported that nearly 60% of American workers have experienced racial and ethnic harassment. A study in Europe indicates that between 10% and 15% of the workforce experience workplace harassment (Zapf et al., 2011).

Workplace harassment can take various forms and mostly based on various group characteristics such as; race, sex, age or color (Raver, Nishii, Kozlowski & Steve, 2010). Workplace bullying has been identified with many serious health problems (Felblinger, 2008). Victims of workplace harassment report more symptoms of depression and anxiety, which have been known to be risk factors of mental and cardiovascular diseases (Hansen, 2006). The World Health Organization reports that more than 300 million people of all ages suffer from depression globally (WHO, 2017). The causes of depression are complex, and research has found several risk factors related to it (Gotlib & Hammen, 2009). Montano, Reeske, Franke, & Huffmeier, (2017), in a meta-analytic study, found that destructive leadership, manifested by aggressive and authoritative attitudes in the workplace is a major cause of depression and poor psychological well-being.

Workplace sexual harassment has been on the rise in the United States in recent times (Levine, 2017). The annual prevalence of depression in 2010 in women and men was 5.5% and 3.2% (Whiteford & Degenhardt, 2010). One factor that might explain the difference in diagnosis of depression between men and women is sexual harassment which mostly occurs at the workplace (Levine, 2017).

Despite the reported rise in workplace harassment in the United States in recent years, there have not been many studies on its relation to depression. Is there any link between workplace harassment and depression? Do women experience more depression due to workplace harassment than men? Does depression has a relationship with marital status? These are the questions this research seeks to answer.

## METHODOLOGY

### Sample

We analyzed data from the National Health Interview Survey (NHIS) in 2015 with total participants (n= 33,672). The NHIS is a survey of civilian non-institutionalized population in the United States conducted by the Centre for Disease Control (CDC). The study design used was cross-sectional with cluster sampling. For our analysis we included all adults who participated in the survey and had continuous employment in the past year. This study relied on secondary data, containing no personal identifiers; therefore, no institutional review board approval was necessary. Contained in the survey were sociodemographic characteristics, habits, healthcare use, physician diagnosed medical conditions and measures of physical health.

### Measures

Variables collected included depression (dependent variable), workplace harassment (independent variable). Confounders such as; demographic characteristics (age, sex, race, marital status) and employment characteristics (income, number of years on the job) of study participants were also collected. Status of depression was determined by asking questions like “does depression cause difficulty with activity or not?” Workplace harassment was determined by asking,” have you ever been harassed on the job?”

### Data analysis

Data was analyzed using IBM Statistical Package for the Social Sciences (SPSS) for windows, version 25.0.

In the primary approach, we conducted descriptive analyses of demographic characteristics and employment characteristics. A Chi-square test was done to assess the relationship between variables. The primary effect of interest was the relationship between workplace harassment and depression. Other variables such as age, sex, race, marital status, income, number of years on the job and workplace safety were included in the analysis and their relationships with depression were assessed. Unadjusted odd ratios (OR) between depression and predictors were calculated using bivariate logistic regression. Multivariate logistic regression was used to determine adjusted OR while adjusting for potential confounders. All values were computed at 95% confidence interval.

## RESULTS

### DESCRIPTIVE ANALYSIS

Sample characteristics are in Table 1. The sample included 33,672 adults with ages ranging from 18 – 65years. The study population was predominantly females (n= 18601, 55.2%) and whites (n= 24230, 79.2%). Majority of the participants were married (n=14787, 52.3%) followed by those who had never married (n= 7708, 27.3%). More than half of the participants were paid per hour ((56%). More than two-thirds (72.1%) of the participants had worked for 1-34years within their current workplace. 6.3% of the participants has experienced depression and 9.9% of the participants had experienced some form of workplace harassment.

**TABLE 1 -.**
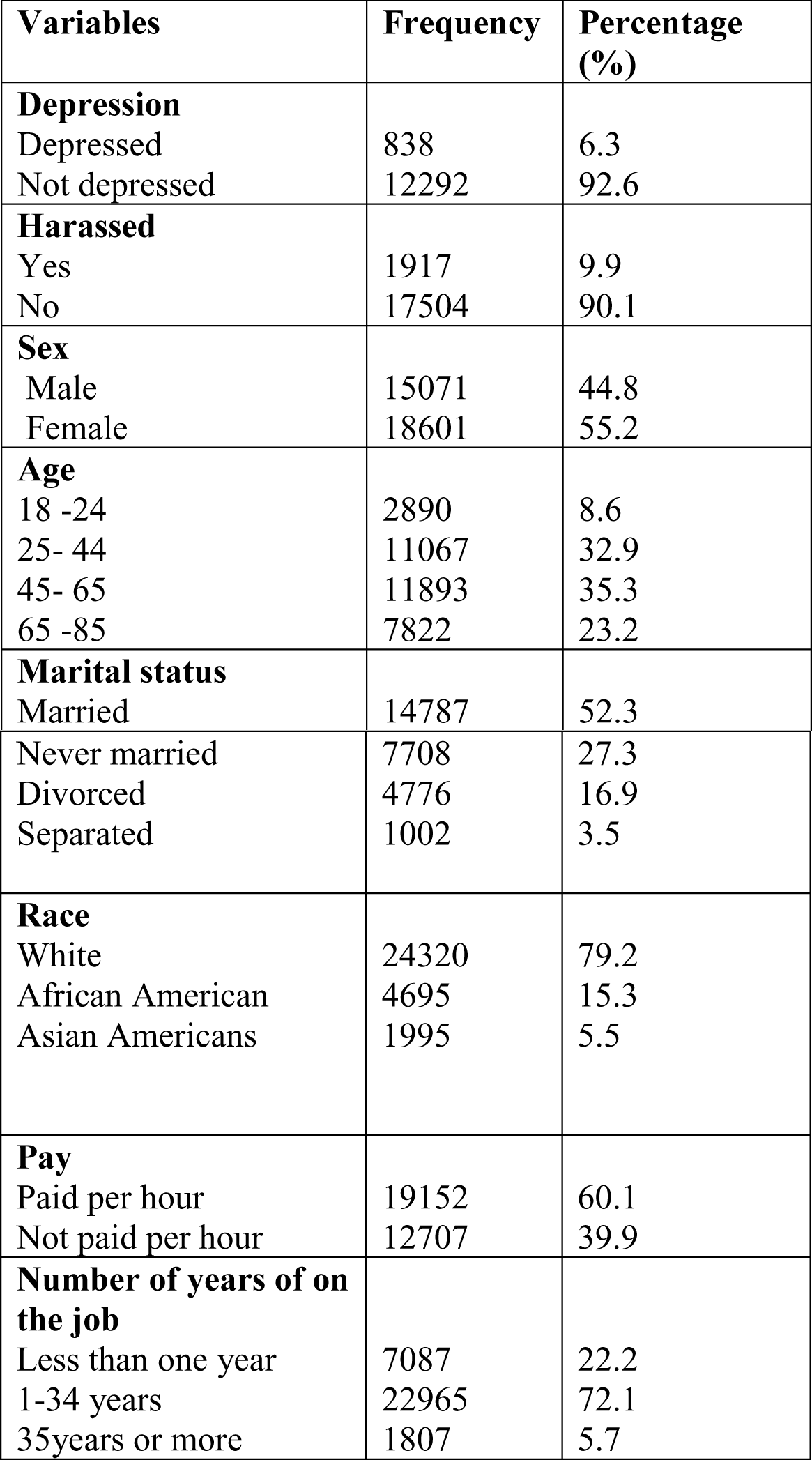
Descriptive analysis of variables of participants (N= 33672)

| **Variables** | **Frequency** | **Percentage (%)** |
| --- | --- | --- |
| **Depression** |  |  |
| Depressed | 838 | 6.3 |
| Not depressed | 12292 | 92.6 |
| **Harassed** |  |  |
| Yes | 1917 | 9.9 |
| No | 17504 | 90.1 |
| **Sex** |  |  |
| Male | 15071 | 44.8 |
| Female | 18601 | 55.2 |
| **Age** |  |  |
| 18 -24 | 2890 | 8.6 |
| 25- 44 | 11067 | 32.9 |
| 45- 65 | 11893 | 35.3 |
| 65 -85 | 7822 | 23.2 |
| **Marital status** |  |  |
| Married | 14787 | 52.3 |
| Never married | 7708 | 27.3 |
| Divorced | 4776 | 16.9 |
| Separated | 1002 | 3.5 |
| **Race** |  |  |
| White | 24320 | 79.2 |
| African American | 4695 | 15.3 |
| Asian Americans | 1995 | 5.5 |
| **Pay** |  |  |
| Paid per hour | 19152 | 60.1 |
| Not paid per hour | 12707 | 39.9 |
| **Number of years of on the job** |  |  |
| Less than one year | 7087 | 22.2 |
| 1-34 years | 22965 | 72.1 |
| 35years or more | 1807 | 5.7 |

### BIVARIATE ANALYSIS

Adults in the study were divided into two groups; depressed (n=838) and not depressed (n=12292). A greater percentage (7.1%) of the adults who were harassed experienced depression as compared to those who were not harassed (4.6%). There was no statistically significant difference in depressive episodes between males and females. The prevalence of depression was highest (14.1% and 10.9%) in people of age groups 18-24years and 25-44years respectively. The lowest rate of depression was seen among adults of age group 65-85years. The highest rate of depression was seen among whites (6.5%). The prevalence of depression was almost equal among African and Asian Americans (5.5% and 5.6%) respectively. Adults who were married had the lowest prevalence of depression (4.6%). The highest rate of depression was seen among participants who had never married and those that were separated 10.0% and 10.9% respectively. Participants who were paid per hour experienced more depression (7.4%) as compared to those who are not paid per hour (4.4%). Workers who had spent less than one year on the job had the highest prevalence of depression (12.8%). The least incidence of depression was seen among participants who had spent 35years or more on job (2.4%). The chi-square tests indicated statistical significance between depressive episodes and all the variables except sex (p-value < 0.05).

### ADJUSTED AND UNADUSTED ODD RATIOS

A linear model regression model was constructed to predict the frequency of depression based on workplace harassment and the demographic variables. Adults who were harassed were 0.39 times more likely to be depressed than those who were not harassed (95%CI: 0.45-0.81), after adjusting for sex, age, race, marital status, pay, and number of years on the job, the odd was almost the same [AOR (95%CI):0.55(0.40-0.77)] (Table 3). There was no statistically significant difference in prevalence of depression between males and females. The odd of been depressed was significantly higher in adults who had never married [OR 2.52(1.83-3.49)] were not paid per hour [OR 1.74(1.48-2.05)] and had worked for more than a year [OR 2.67 (2.29-3.13)] than their counterparts. The odds of been depressed was significantly decreased by 0.84 (C.I. – 0.12-0.22) and 0.67(0.27-0.40) across age groups when compared to age category 18-24.

**TABLE 2 –.** Relationship between participants with depression and those without depression (N= 33672)

| **VARIABLES** | **DEP + (%)** | **DEP – (%)** | **P – VALUE** |
| --- | --- | --- | --- |
| **Harassed** |  |  |  |
| Yes | 42(7.1%) | 552 (92.9) | 0.01 |
| No | 193(4.6%) | 4021(95.4) |  |
| **Sex** |  |  |  |
| Male | 320(6.3%) | 4747(93.7%) | 0.80 |
| Female | 518(6.4%) | 7545(93.6) |  |
| **Age(yes)** |  |  |  |
| 18 -24 | 56 (14.1) | 333 (84.1) | 0.01 |
| 25- 44 | 257(10.9) | 2074(87.8) |  |
| 45- 65 | 390 (7.4) | 4823 (91.6) |  |
| 65 -85 | 135 (2.6) | 5062 (94.6) |  |
| **Race** |  |  |  |
| White | 652 (6.5) | 9270 (92.6) | 0.01 |
| Black/ African American | 108 (5.5) | 1831 (93.2) |  |
| Asian Americans | 27 (5.6) | 443 (91.7) |  |
| **Marital status** |  |  |  |
| Married | 245 (4.6) | 4975 (94.2) | 0.01 |
| Never married | 216(10.0) | 1908 (88.7) |  |
| Divorced | 196 (8.2) | 2164 (90.9) |  |
| Separated | 49 (10.9) | 398 (92.1) |  |
| **Pay** |  |  |  |
| Paid per hour | 576(7.4) | 7156(92.6) | 0.01 |
| Not paid per hour | 204(4.4) | 4330(95.6) |  |
| **Number of years of on the job** |  |  | 0.01 |
| Less than one year | 267(12.8) | 1799 (86.4) |  |
| 1-34 years | 489 (5.2) | 8789 (93.7) |  |
| 35years or more | 24 (2.4) | 971 (96.0) |  |

**TABLE 3 –.**
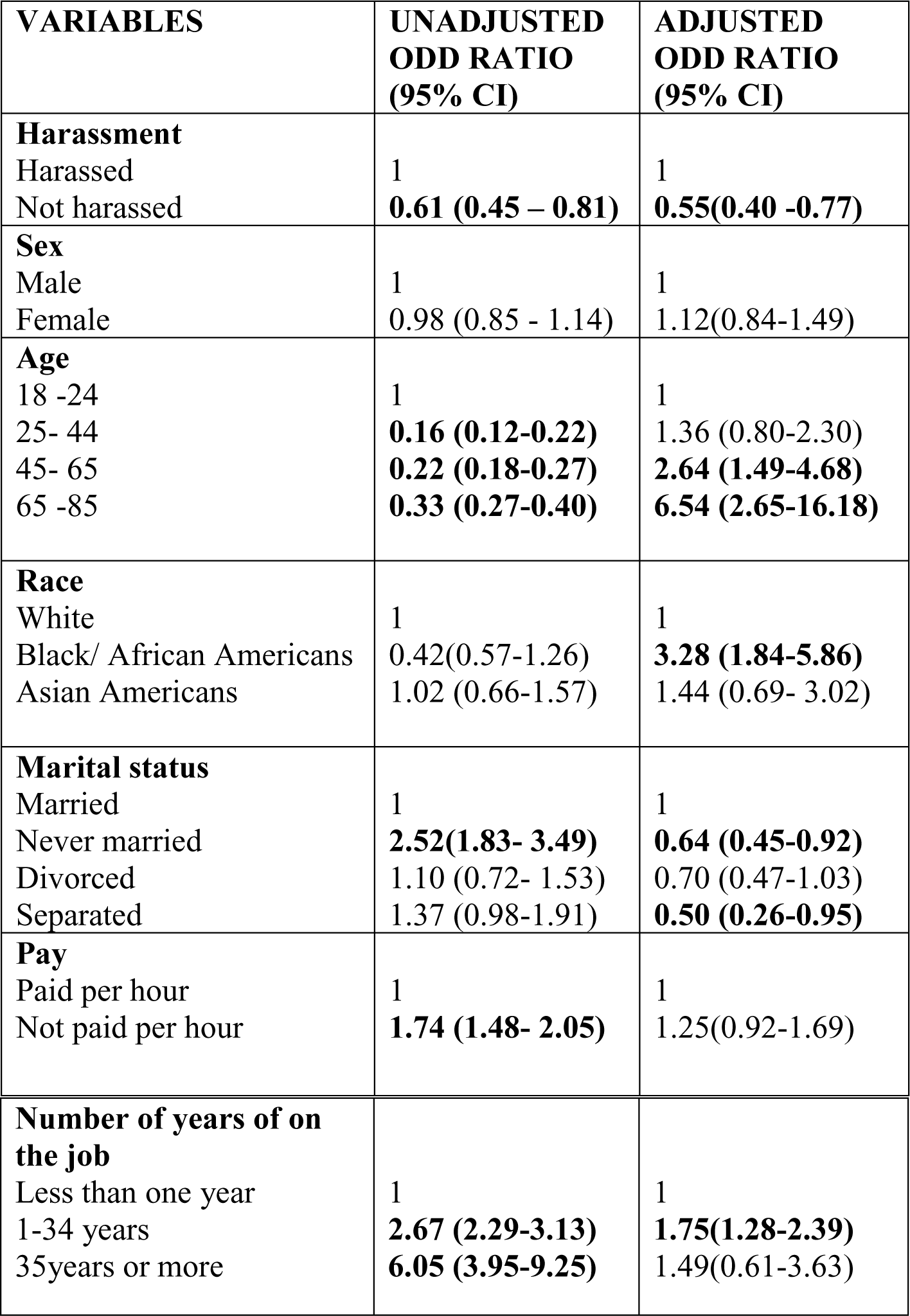
ADJUSTED AND UNADJUSTED RATIOS.

| **VARIABLES** | **UNADJUSTED**<br>**ODD RATIO**<br>**(95% CI)** | **ADJUSTED**<br>**ODD RATIO**<br>**(95% CI)** |
| --- | --- | --- |
| **Harassment** |  |  |
| Harassed | 1 | 1 |
| Not harassed | **0.61 (0.45 – 0.81)** | **0.55(0.40 -0.77)** |
| **Sex** |  |  |
| Male | 1 | 1 |
| Female | 0.98 (0.85 - 1.14) | 1.12(0.84-1.49) |
| **Age** |  |  |
| 18 -24 | 1 | 1 |
| 25- 44 | **0.16 (0.12-0.22)** | 1.36 (0.80-2.30) |
| 45- 65 | **0.22 (0.18-0.27)** | **2.64 (1.49-4.68)** |
| 65 -85 | **0.33 (0.27-0.40)** | **6.54 (2.65-16.18)** |
| **Race** |  |  |
| White | 1 | 1 |
| Black/ African Americans | 0.42(0.57-1.26) | **3.28 (1.84-5.86)** |
| Asian Americans | 1.02 (0.66-1.57) | 1.44 (0.69- 3.02) |
| **Marital status** |  |  |
| Married | 1 | 1 |
| Never married | **2.52(1.83- 3.49)** | **0.64 (0.45-0.92)** |
| Divorced | 1.10 (0.72- 1.53) | 0.70 (0.47-1.03) |
| Separated | 1.37 (0.98-1.91) | **0.50 (0.26-0.95)** |
| **Pay** |  |  |
| Paid per hour | 1 | 1 |
| Not paid per hour | **1.74 (1.48- 2.05)** | 1.25(0.92-1.69) |

| **Number of years of on the job** |  |  |
| --- | --- | --- |
| Less than one year | 1 | 1 |
| 1-34 years | **2.67 (2.29-3.13)** | **1.75(1.28-2.39)** |
| 35years or more | **6.05 (3.95-9.25)** | 1.49(0.61-3.63) |

Adjusted for all other variables, the odds of depression was significantly higher in age group 65-85 [AOR (95%CI):6.54(2.65-16.18)] (Table 3), African Americans [AOR (95%CI):3.28(1.84-5.86)] (Table 3), and workers who had worked for 1-34years [AOR (95%CI):1.75(1.28-2.39)] (Table 3) than their counterparts.

## DISCUSSION

The study found that workplace harassment is positively related to depression. Age was found to have the highest overall effect on depressive episodes whiles marital status had the lowest. The single highest predictor was adults who had spent more than 35years on the job. The finding is consistent with the results from a study done by Verkuil, Atasayi & Molendijk, (2015). Another study by Khuchandani & Price (2014) found a positive correlation between workplace harassment and risk depression, obesity and cardiovascular diseases.

The observed association can be explained by the stress model that states that long periods of stress can negatively affect somatic as well as mental health (Kivimaki at al., 2003). Workplace harassment can be considered as source of stress that affects emotional well-being, by causing changes in the neuroendocrine, autonomic and immune functioning (Kivimaki et al., 2003).

There was also a correlation between marital status and depression. Adults who had never married or were out of marriage were seen as more likely to be depressed than those who were married. This is consistent with the findings by Simon, (2002), who found an association between marital status and depression.

### Limitations and future research

The study demonstrated statistically significant relationship between workplace harassment and depression. However, a variety of factors may have imposed limitations on the results. First, the use of self-reported data has the potential of introducing recall bias and social desirability.

Second, the use of cross-sectional study design does not allow us to make conclusions of a causal nature. Another limitation of this study is that the variables such as depression and harassment was not indicative of participant’s present situation, whether they were currently been harassed or not. This made establishing ongoing relationship between workplace harassment and depression very difficult.

We suggest that future research should consider the interaction between various forms of workplace harassment and ways to limit its incidence.

## Conclusion

Existing research makes clear that inappropriate behavior in the workplace is rampant in many organizations. Workplace harassment may consist of: managers towards employees or their fellow managers, employees towards their fellow employees or managers, or the organization itself. Several forms of cations and inactions can be termed as workplace harassment. Sexual harassment, physical abuse, work sabotage, name calling, eye rolling, dirty looks and teasing may all be termed as workplace bullying. Organizations must move quickly to address bullying and create conducive working environments for their employees, since the legal and economic consequences of failing to address workplace harassment are huge.

This study used data from the National Health Interview Survey 2015 to figure out the relationship between workplace harassment and depression. We found a positive correlation between the said variables though other variables such as; age, race and marital status can influence depression.

